# Open Source LED controller for circadian experiments

**DOI:** 10.1101/201087

**Authors:** Maite Ogueta, Luis Garcia Rodriguez

## Abstract

Controlling the light conditions when performing a circadian experiment is a key factor. In this article a simple, affordable and open source LED controller is presented. Using commercial available solutions like and Arduino [1] and LED stripes the cost is dramatically reduced.

## Introduction

Since the 1990s, many laboratories have focused their studies on the circadian clock, using many different model organisms. The study of circadian rhythms requires a controlled environment where light and temperature can be modified to study the different features of the clock. This is usually done within commercially available incubators (e.g. Percival Scientific Inc.), that allow the user to control some of the parameters (lights on-off, temperature cycles with ramping..). But so far these incubators can not be programmed to change automatically from one program to another, and the settings can not be adapted if something different is needed. There are other commercially available solutions to control and program the lights, such as the HLT Control V2.0 software in combination with an LED light box (Hoenig Lichttechnik, Bonstetten, Germany) [4]. Other labs have even developed home made systems in order to have a precise control of the lightning conditions [5], [2].

This work present an LED controller that is independent of the incubator type, shape and size, without requiring an expert level of electronics or programming. Its modular design allows a wide range of personal configurations. The LED strip can be changed at any moment, allowing an easy way to use lights of different wavelengths. Due to its open source nature, the code can be modified and adapted to implement new functionalities or to adjust the existent ones. The present version of the software allows not only a rectangular light dark cycles, but has also a ramping mode to simulate dawn and dusk, and a pulsing mode, useful for experiments such as optogenetics. It can also be programmed to change from one mode to another after a desired time.

In this article basic information of the hardware and software are provided, and the code is made available.

## 1. Hardware

An overall view of the different parts of this LED controller is provided in figure 1. There is a Computer that controls one or several Arduinos. Each Arduino has a shield where the LED strip and the power supply are attached (Fig. 2).

**Figure 1.**
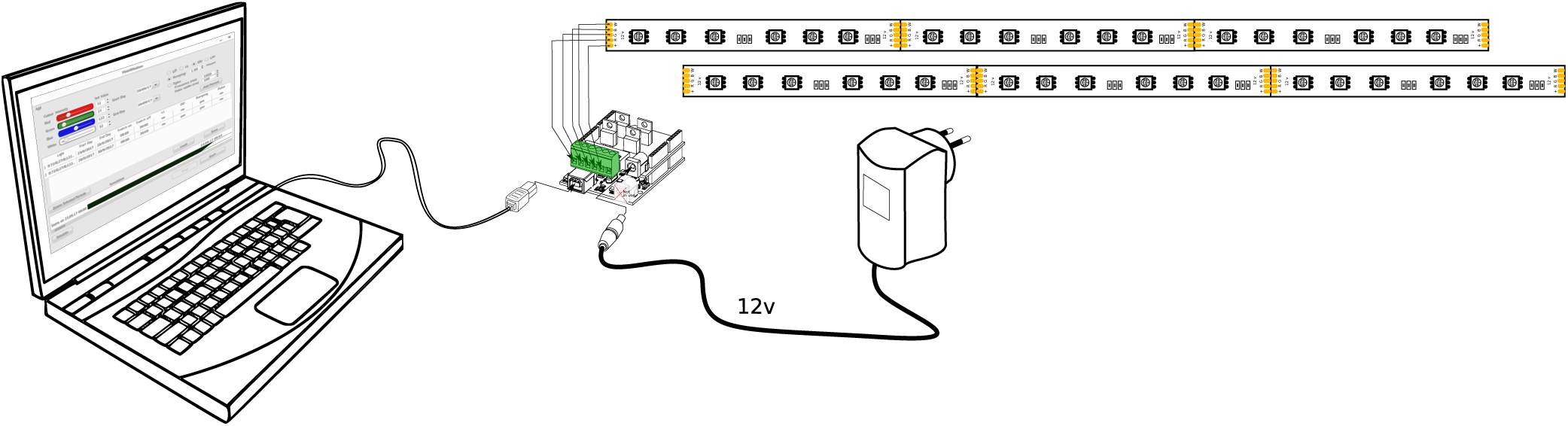
A general view of how all the components are connected. The laptop computer run the Desktop App (led-control). The Arduino with the shield is connected to the computer by USB and to the 12v power supply. The LED strip is connected to the Shield by means of the green hub.

**Figure 2.**
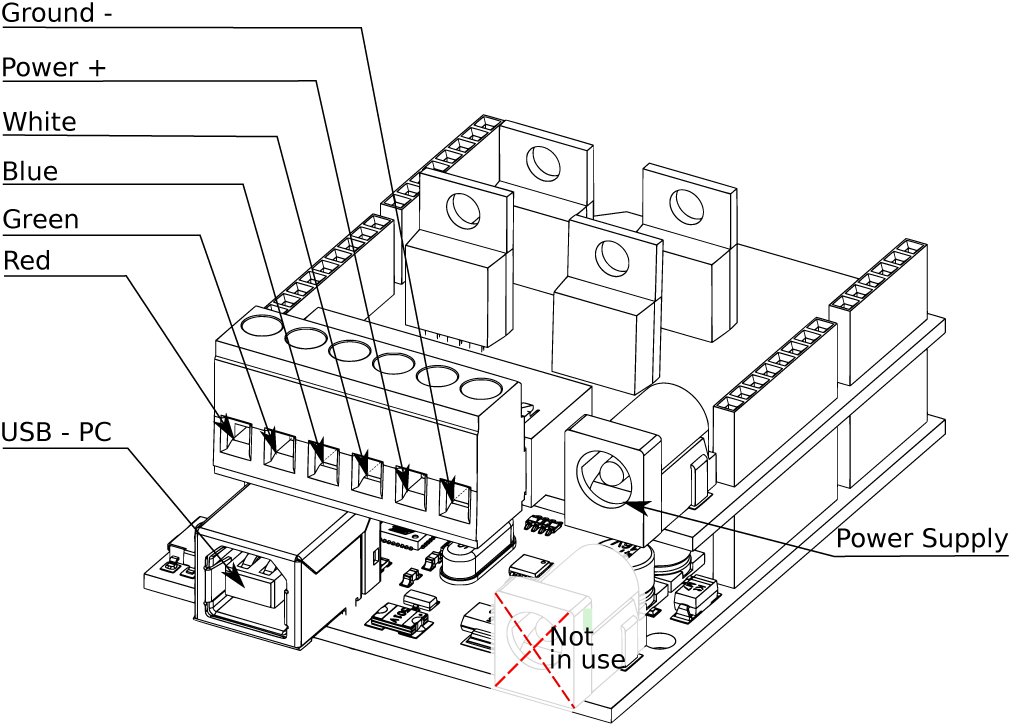
Arduino + Arduino Shield.

### 1.1 Arduino

The hardware is based in an Arduino microcontroller expanded with a custom made shield (see Figure 2). This shield is a custom made PCB that provides the Arduino with the capabilities to switch on and off lights (or other devices if desired) up to 5A and 12V. Blueprints to produce this PCB can be found in the github repository [3]. It is also possible to find commercially available shields that provide similar capabilities to the Arduino (i.e Arduino Motor Shield rev3 https://store.arduino.cc/arduino-motor-shield-rev3), but in that case the wiring and the Arduino Firmware (section 2.2) will change. With some basic knowledge of electronics, it is possible to amend it to make it work.

### 1.2 LEDs

The LED strip can be any commercially available 12V LED strip. The overall length used depends on the consumption of the LED strip, with a maximum voltage allowed by the shield of 60W. Individual LEDs and arrays of LED’s also can be used.

The software and the Arduino shield, as they are designed now, support up to 4 independent channels. A RGBW (red, green, blue, white) LED strip was used for the original design, but any other LED combination that matches the specifications of the Arduino Shield can be used.

The LED strips are connected to the shield following the scheme showed in figure 2, with each of the channels connected to a different port, plus the power (or +) and the ground (or -). This will depend on the type of LEDs that are used. LED strips with other colours like violet, amber, orange or yellow can still be used with the new color connected to any of the 4 ports, although the software will not automatically reflect this change. That means that if yellow LEDs are connected to red port, they will be operated with the red signal in the software.

Individual LEDs can be used if in series resistor is provided, but a minimum knowledge in electronics is required to calculate the right value of the resistor.

### 1.3 Power supply

The power supply provides energy for the LEDs and it is connected to the shield in the power supply port (see Figure 2). It is worth mentioning that the original power supply port of the Arduino is not used. Connection of the power supply in this port can damage the Arduino board.

The specifications of the connector are: FC68148 DC10A dc power connector 5A, 12v, 2.1mm center negative. Full characteristics can be found in the datasheet (Farnell datasheet http://www.farnell.com/datasheets/317160.pdf).

The specifications of the power supply are dependent on the LED strip selected. The maximum specifications supported by the shield are 12v and 5A (60W). This limitation comes from the dc power connector. However, it could be possible to change this for another type of connector.

### 1.4 Computer

While the Arduino and the shield could be considered the actuator or worker, the computer is the brain of the system. It runs the desktop app (or program) that sends the right commands to operate the Arduino.

The minimum specifications for this computer is an Intel i3 or ARM cortex-A53 processor with 1 GB ram. It can also run in inexpensive micro computers like Rapsberry pi. The recommended model is number 3 (last model in the moment of writing). Multiple Arduinos can be connected to a single computer with a USB hub, thus reducing the overall costs of the installation. The recommended operating system for this computer is Linux, for example Ubuntu, Fedora linux or elementary OS. This will simplify the installation of the software, without the need of software licences.

## 2. Software

The software used in this project has been custom made, and it has been developed with the circadian requirements in mind.

It consists of two separate parts: The Desktop app and the Arduino Firmware. Both of them can be found in the Github repository [3]

### 2.1 Desktop App

The main program runs in the computer specified above (Section 1.4). The program can be started more than once, for each instance (or run) the computer controls one Arduino. In figure 3 some screen shots of the program are presented. The main Window (shown in figure 3 a) is divided in three different parts:

1. Period parameters
2. Period list
3. Simulation and visualization of set of periods.

**Figure 3.**
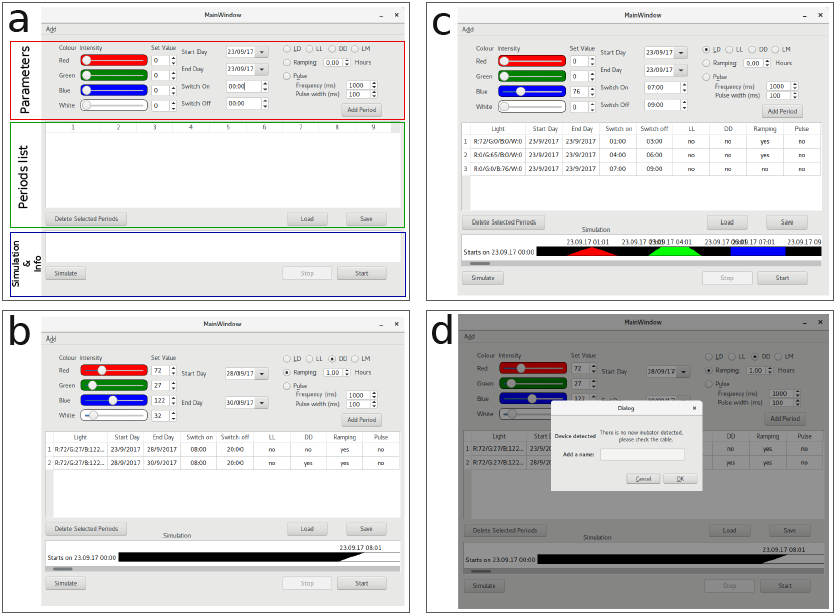
(a). Main view of the program as it starts. (b) Main view after two periods has been added and simulation buttons has been clicked. (c) View showing how periods can be added in sequence to the same day and with different colours. Simulation area shows the color of the period. Variations in ramping are also show: 1 hour ramping for red light, 30 min ramping for green light and no ramping for blue light. (d) Shows Dialog for adding new incubator, the text show is in case no new Arduino controller is attached. The name of the new detected controller needs to be given in the “Add a name” text box.

In part one (figure 3 a, red pannel) the user specifies the parameters to create a period. A period consist on:

- Colour of the light (single or multichannel)
- Intensity of the light (a value between 0 and 255)
- Date when the period starts
- Date when the period finishes
- Hour when lights should go on in this period
- Hour when lights should go off in this period
- Type of period: light:dark (LD), constant darkness (DD), constant light (LL) or light:moonlight (LM)
- Selection of ramping and duration of it (if needed)
- Option to select a pulsing light and its duration.

A set of periods will be further considered as as an experiment.

In the second section (figure 3 a, green pannel) the user can see a list of all the added periods with the actual configuration. In this panel some actions are possible, such us delete periods, save the list for later use, or load a list from a previously saved experiment. This files can be interchanged with colleagues so they can replicate the same experiment. If one period is selected the parameters can be modified in the parameter section and updated them by clicking the button “Update Period”. The periods can be updated in real time and while the experiment is running. The new conditions can take up to one minute to be updated inside the incubator.

More information about usage is presented in the section 2.1.2.

The third and the last section shows a representation of the actual experiment (figure 3 a, blue pannel). By clicking on simulate, a whole simulation is performed with the data provided in the periods, allowing the user to identify possible mistakes or just to check that the program will perform as expected. During the actual experiment, this section will show an actual state of the system. (figure 3 c)

#### 2.1.1 Installation

For an Installation guide please refer to the README in the Github repository [3].

#### 2.1.2 Minimum user manual

Once the program has started the Main Window will be visible (figure 3 a) with an empty period list.

##### Adding new incubators

The first thing to do is to add a new incubator where the LEDs are installed. These are the steps:

1. Connect the USB cable from the arduino in one available USB in the computer.
2. In the program, click “add” in the top left menu.
3. In the prompted window write a new name to identify the new detected Arduino controller and click OK.
4. If in the previous step it is not possible to add a new name and the text shown is the one in figure 3 d, then check the USB connection and the cable and try again.

##### Adding periods

Using the Parameters section (figure 3 a, red area) a new period can be created. Start by selecting the desired colour, each slider will set the intensity of the respective colour, being 0 off and 255 the maximum intensity. The value can be introduced by moving the slider, manually in the value box or by steps with the up and down arrows. By default this colours are mapped with the port of the same name in figure 2. Once the light is selected, the next step is to introduce the start and end dates, in combination with the Switch on and Switch off times that will define how the period is going to perform. The period can be as short as 1 minute and as long as desired. In figure 3 c, three periods for the same day with different colours and intensities are shown.

Next step is to select the type of period from one of the following:

- LD: Light/Dark (default)
- LL: Light/Light (selected light will be on for 24 hours between selected dates)
- DD: Light/Dark (lights will be off the 24 hours between selected dates)
- LM: Light/Moon (this will keep a really small amount of white light during the night, this is a beta feature and should be tested before starting an actual experiment)

The last step defines some other characteristics of the light. For example, if the transition between light/dark and dark/light has to be ramped. In this case select the Ramping function. The “Hours” box will define how long this ramping will last. If “Pulse” is selected, then the lights will be switched on and off (pulsed) according to the following parameters: “Frequency” is how often the pulse loop is to be repeated (given in milliseconds) and “Pulse Width” is the duration of the light pulse for each loop.(i.e for 1000ms Frequency and 100ms Pulse width, every second the light will be on for 0.1 seconds).

Once the period is ready by clicking “Add Period” it will appear in the Period list (figure 3 a, green area).

This can be repeated until the desired number of periods is entered.

##### Update periods

As stated before, periods can be updated by selecting one period in the Period list and then changing the parameters as shown in the previous section. The action can be cancelled at any moment clicking “Cancel”. For the period to take effect “Update period” button has to be clicked. Changes will be reflected immediately in the Period list and if the experiment is already running they will also change the experiment conditions (this can take up to a minute).

##### Save and load experiments

Once the period list is completed that will be considered an experiment and there is the option to save it for later use. By clicking “Save” a system dialogue will appear asking where to save the experiment.

Previously saved experiments can be loaded by using “Load” button. The loaded experiment can be updated as well.

##### Simulate experiment

One way to check if you have programmed the experiment correctly is to use the “Simulate” button. This will run a simulation for the duration of the whole experiment. Examples of simulations can be seeing in figure 3 b and c.

##### Start and Stop experiment

To start the experiment, the “start” button should be clicked. This will prompt a window allowing the user to select one of the available incubators (previously added Arduinos). If no Incubators are available, that means that all the Arduino controllers are in use.

To stop the experiment is really important to do it by using the “Stop” button. This will make the attached Arduino controller available again to run another experiment. If that is not the case, there are chances that the Arduino controller will not be released and get corrupted, until the config.cfg file is removed (The file can be found in the same folder as the source code). This will imply that you have to re-add all the Arduino controllers again.

### 2.2 Arduino Firmware

In order to receive the commands from the main program and correctly switch on and off the lights, the Arduino needs to run a small program that manages the communication with the computer and the outputs for the lights. This small program is called firmware‥

To load this firmware into an Arduino you can follow the instructions from the official arduino website [1].

This Firmware uses serial communication to receive orders. In case the user wants to write his/her own desktop program to control the lights, here are some basic information.

Arduino is hearing constantly over serial waiting for specific commands to arrive. The basic structure of this commands is as follows:

1. A character: R,G,B,W,F,N or C
2. An integer (number)

To switch on red (R), green (G), blue (B) and White (W) LEDs the command is the respective letter followed by an integer between 0 and 255. The number will indicate the intensity of the light between 0% = 0 and 100% = 255. For example:

- “R25”: Red light set to 25.
- “R25G25B255”: Red light set to 25, green to 25 and blue to 255.
- “B0”: Blue light off.

To create a pulse of light instead of a continuous light value, it is necessary to use the Frequency command (F). It consists of the letter followed by two integers, this two integers are frecuency of the pulse, in milliseconds, and width of the pulse, also in milliseconds. For example: “F1000/100” will create a pulse of lights on for 100 milliseconds every 1 second (1000ms)

Sending the letter N (No pulse) without any integer will stop the pulses, i.e. N cancels the effect of F.

Sending letter C (Communicate), will force the Arduino to respond with “Led Controller”. This command is useful for device discovery, and it is used internally to detect and add new devices in the main program.

## Acknowledgments

This development has been possible thank to Ralf Stanewsky that has provided with the initial founding. He also was part of the design process providing valuable knowledge and has helped to establish the basic requirements. We would like to acknowledge also Adam Bradlaugh, who suggested the addition of the pulsing function and has been a tester of the program.

